# Mechanistic Elucidation of CLIC1 Membrane Insertion via Structural and Dynamic Modulation

**DOI:** 10.1101/2025.01.19.633805

**Authors:** Joseph Cassar, Angela Serrano-Sanchez, Jack Bragg, Encarnacion Medina-Carmona, Felipe Ossa, Gary Thompson, Vahitha Abdul-Salam, Lorena Varela Alvarez, Jose L. Ortega-Roldan

## Abstract

Chloride Intracellular Channel 1 (CLIC1) is a metamorphic protein capable of transitioning from a soluble cytoplasmic state to a membrane-bound chloride channel. This conformational shift, crucial for physiological processes such as cell volume regulation, electrical excitability, and angiogenesis, is linked to pathological conditions including malignancies and cardiovascular diseases. Despite its significance, the molecular mechanism driving CLIC1’s membrane insertion has remained elusive.

Using an integrated structural biology approach combining NMR spectroscopy, SAXS, biophysical methods, and mutagenesis, we uncover the dynamic landscape underpinning CLIC1 function. Solution NMR and SAXS reveal that CLIC1 adopts a conformational ensemble in equilibrium, characterized by a compact ground state and a partially extended state exposing key membrane-interacting regions. Zn^2+^ binding acts as a critical trigger, inducing structural rearrangements, increasing protein flexibility, and promoting oligomerization essential for membrane insertion.

Our findings demonstrate that structural flexibility, particularly within dynamic loop regions and interdomain linkers, is intrinsic to CLIC1’s ability to adapt to membrane interactions. Zn^2+^-induced dimerization and tetramerization were identified as key steps preceding insertion, with mutations in the transmembrane (TM) region revealing pivotal roles for residues R29 and W35 in modulating protein dynamics, oligomerization and insertion efficiency.

This study provides a mechanistic framework for CLIC1’s transition to its membrane-bound state, offering insights into the interplay between conformational dynamics, oligomerization, and metal ion modulation. These findings pave the way for targeted strategies to regulate CLIC1 activity in pathological conditions, underscoring its potential as a therapeutic target.

## INTRODUCTION

Chloride Intracellular Channel family consists of a group of highly homologous proteins with a striking feature, their ability to change their structure upon activation from a soluble form into a membrane-bound chloride channel, translocating from the cytoplasm to intracellular membranes(Tulk et al., 2000; Valenzuela et al., 1997). CLIC1 is the best characterised of the CLIC protein family and seems to act as a chloride channel in its membrane-bound form. It is expressed intracellularly in various cell types, especially abundant in the heart and skeletal muscle(Ulmasov et al., 2007). CLIC1 has been implicated in the regulation of cell volume,(Singh et al., 2007) electrical excitability,(Averaimo et al., 2014) diNerentiation,(Wang et al., 2011) cell cycle(Valenzuela et al., 2000) and cell growth and proliferation.(Tung & Kitajewski, 2010) DiNerent studies have reported high CLIC1 expression in a range of malignant tumours(Gritti et al., 2014; Zhang et al., 2015) and cardiovascular diseases such as pulmonary hypertension(Abdul-Salam et al., 2010) and ischaemic cardiomyopathy.(Gronich et al., 2010) Only the channel form of CLIC1 has been shown to have pro-proliferative and angiogenic activity.(Carlini et al., 2020; Xu et al., 2016) Therefore, understanding the mechanism of CLIC1 membrane insertion is crucial, as specific inhibition of CLIC1 translocation has the potential to reverse endothelial injury and prevent subsequent vascular disorders.

The membrane translocation mechanism of CLIC1 has remained enigmatic for over 30 years. The initial hypothesis by Littler et al (Littler et al., 2004) was based on oxidation. This theory, however, has been disputed by others since.(Hossain et al., 2016, 2017; Jiang et al., 2012; Stoychev et al., 2009; Tulk et al., 2002) Our lab recently discovered that CLIC1 binds to Zn^2+^, triggering its membrane insertion(Varela et al., 2022) and demonstrated a reproducible two-step mechanism whereby upon Zn^2+^ binding, CLIC1 inserts in the membrane and subsequently activates chloride eNlux upon acidification

The CLIC1 crystal structure reveals a compact, globular fold incompatible with the large conformational changes required for membrane insertion. To address this paradox, we employed solution NMR, SAXS, and other biophysical tools to investigate CLIC1’s dynamic behaviour in solution and its structural rearrangements induced by Zn^2+^. By integrating these approaches, we provide a comprehensive mechanistic model for CLIC1’s membrane insertion and channel formation.

## MATERIALS AND METHODS

### Protein Expression and Purification

Recombinant CLIC1 was expressed in Escherichia coli C43 cells using LB or labelled M9 minimal media and purified using Ni-NTA affinity chromatography followed by size-exclusion chromatography (REF). Constructs were cleaved with TEV protease or uncleaved to evaluate the influence an N-terminal His-tag on oligomerization.

### NMR Experiments

Clic1 NMR samples were collected at 30°C on the Bruker Avance III spectrophotometer with a 1H frequency of 600MHz using a QCIP cryoprobe. Samples in a 50mM NaCl 50mM Hepes pH 7.4 buffer were supplemented with 5% D20 and placed in a 5mm Shigemi tube. All spectra were processed in NMRPipe and viewed and analysed using NMRviewJ (Delaglio et al., 1995; Johnson, 2004).

Backbone assignments were collected using standard TROSY-based triple-resonance experiments on a Bruker Avance III HD with a 1H frequency of 800mHz and a TCI cryoprobe.

TROSY-based15N T1, T2, heteronuclear NOE and relaxation dispersion experiments were collected at 600MHz in the absence of Zn2+ and in the presence of a molar ratio CLIC1:Zn2+ of 0.5.

NH Residual Dipolar Couplings (RDCs) were collected in 8% PEG/Hexanol. The D_2_0 splitting of the sample was 23Hz. A TROSY-HSQC and a standard HSQC pair was collected in aligned and unaligned samples to calculate the RDC values.

CLIC1 titrations were performed on 300µM cleaved and uncleaved CLIC1 samples upon addition of ZnSO_4_. Final Zn^2+^ concentrations of 0.25MR, 0.5MR and 1MR were used and to the final 1MR zinc sample, 1mM EDTA was added.

### SAXS Data Collection and Analysis

Samples were collected on Beamline 21 in the synchrotron at Diamond light source. Samples existed between 10-20mg/ml and were in a 50mM Hepes buffer pH 7.4 with 5% glycerol. SEC-SAXS was implemented with a KW-403 Shodex SEC column. SEC-SAXS data was processed and analysed using the ATSAS and Scatter packages(Manalastas-Cantos et al., 2021; Rambo, 2015).

DAMMIF(Petoukhov et al., 2012) was used to provide a visual aid of the average structure of CLIC1. The FOX server(Schneidman-Duhovny et al., 2016) was used to compare the experimental scattering data against the predicted scattering of structural models.

The structures contributing to the ensemble in solution were calculated using EOM(Petoukhov et al., 2012). CLIC1 was split into 4 domains to allow movement of all 3 loops identified via NMR to be flexible and to replicate the dynamics of CLIC1 seen in the NMR. The residues allowed to reorientate were 35-45, 90-100 and 150-165, which the rest of the sequence acting as rigid bodies.

### Mass Photometry

10µL of the CLIC1/nanodisc prepared as described elsewhere(Medina-Carmona et al., 2020) was applied to 10 µL buffer on a cover slip resulting in a final concertation of 100nM. The data was collected on a Refeyn OneMP (Refeyn Ltd, UK) mass photometry system. Movies were acquired for 60 seconds. The mass was calculated using a standard protein calibration curve.

## RESULTS

### NMR Reveals Flexible Regions in CLIC1

Backbone resonance assignment of CLIC1 identified 187 out of 227 non-proline residues, with missing resonances corresponding to the putative TM helix and adjacent C-terminal helix (Figures S1-2). 15N relaxation and relaxation dispersion studies revealed three highly dynamic regions (Figure 1A-C): 1- **Foot Loop (140–170):** Fast nanosecond to microsecond dynamics, consistent with the absence of electron density in crystal structures. 2 - **Interdomain Loop (80–100):** Elevated R2/R1 ratios and chemical exchange at residues 88, 89, 92, and 98, indicating slow, microsecond to millisecond timescale motions. 3- **TM/Loop (35–50):** Differential dynamics across this region suggest distinct structural roles for residues proximal and distal to the TM helix. These findings suggest that structural flexibility is intrinsic to CLIC1, particularly in regions critical for membrane interaction.

**Figure 1.**
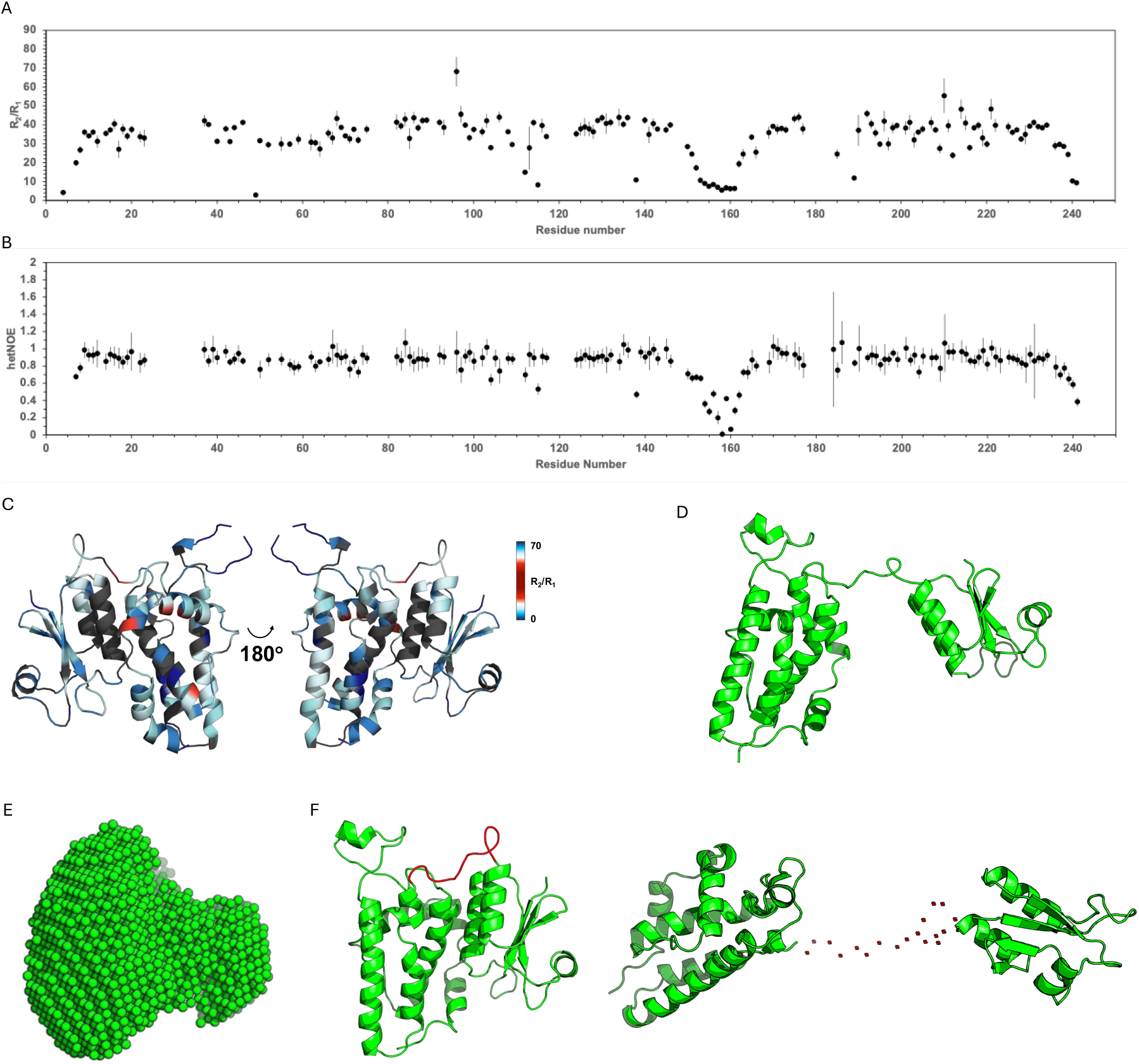
CLIC1 conformational dynamics. A – R2/R1 values of CLIC1 obtained at 600 MHz. B – Heteronuclear NOe values of CLIC1. C – CLIC1 monomeric X-Ray structure (PDB 1k0n) showing dynamic areas. Areas showing fast ps-ns timescale dynamics are represented in dark and light blue. Areas showing slow ms-ns timescale dynamics are shown in red. D – RDC reFined structural model of the average conformation of CLIC1 in solution. E – Beads model of CLIC1 monomer generated using SAXS data in DAMAVER. F - Population ensemble of CLIC1 in solution calculated by EOM. CLIC1 was split into 4 domains to allow movement of all 3 Flexible loops identiFied by NMR. The structure in the left represents the closed state (90%). The structure on the right represent the fully open conformation (10%)., where the N- and C-terminal domains are fully detached.

### NMR RDCs and SAXS Identifies Conformational Ensembles in Solution

Comparison of RDC values, which report on all conformations available in solution, measured on a monomeric CLIC1 construct for 91 non-overlapped, no-flexible residues measured NH RDCs to the predicted RDC values from the X-Ray structure showed a poor fit seen in figure S3. Using the software Module2 we calculated a separate alignment tensor for the N and C terminal domains, due to the flexible linker separating them, using all RDC values in each of these domains respectively (Dosset *et al*., 2001). Our new model of CLIC1 in solution represents an average of all monomeric states seen in solution and differs from the crystal structure of CLIC1 -1K0N with flexibility of the linker bridging the N and C terminal domain (Figure 1D).

SEC-SAXS experiments corroborated that the CLIC1 monomer does not adopt a single conformation in solution. The scattering profile for CLIC1 monomer showed a chi-squared value of 3.86 when compared with the 1K0M crystal structure, indicating significant deviations. P(r) distribution plots supports the existence of a flexible, partially extended conformation, confirming that CLIC1 exhibits significant flexibility, a characteristic essential for its metamorphic nature (Figure 1E and S4).

EOM modelling of the monomer revealed that CLIC1 exists in a dynamic equilibrium between a major compact state (90% population) resembling the crystal structure and a minor elongated state (10%) (Figure 1F and S5). The elongated conformation arises from the separation of the N- and C-terminal domains via the dynamic interdomain loop, exposing the TM region to solvent.

### CLIC1 oligomerization is required for membrane insertion

Size exclusion chromatography (SEC) experiments showed that CLIC1 exists as a mixture between different oligomeric states including monomer, dimer and tetramers, with the monomeric as the main state (Figure 2A and S6). Extension of the N-terminus with a Histidine tag removed the presence of any non-monomer peak, as well as the presence of elongated species in the SAXS curves (Figure S4). CLIC1 membrane insertion has been found to be triggered by Zn^2+^ binding(Varela et al., 2022). Incubation of CLIC1 in the absence of lipids promote oligomerization, shifting the monomer peak to a dimer and tetramer form, and eventually inducing protein aggregation (Figure 2A).

**Figure 2.**
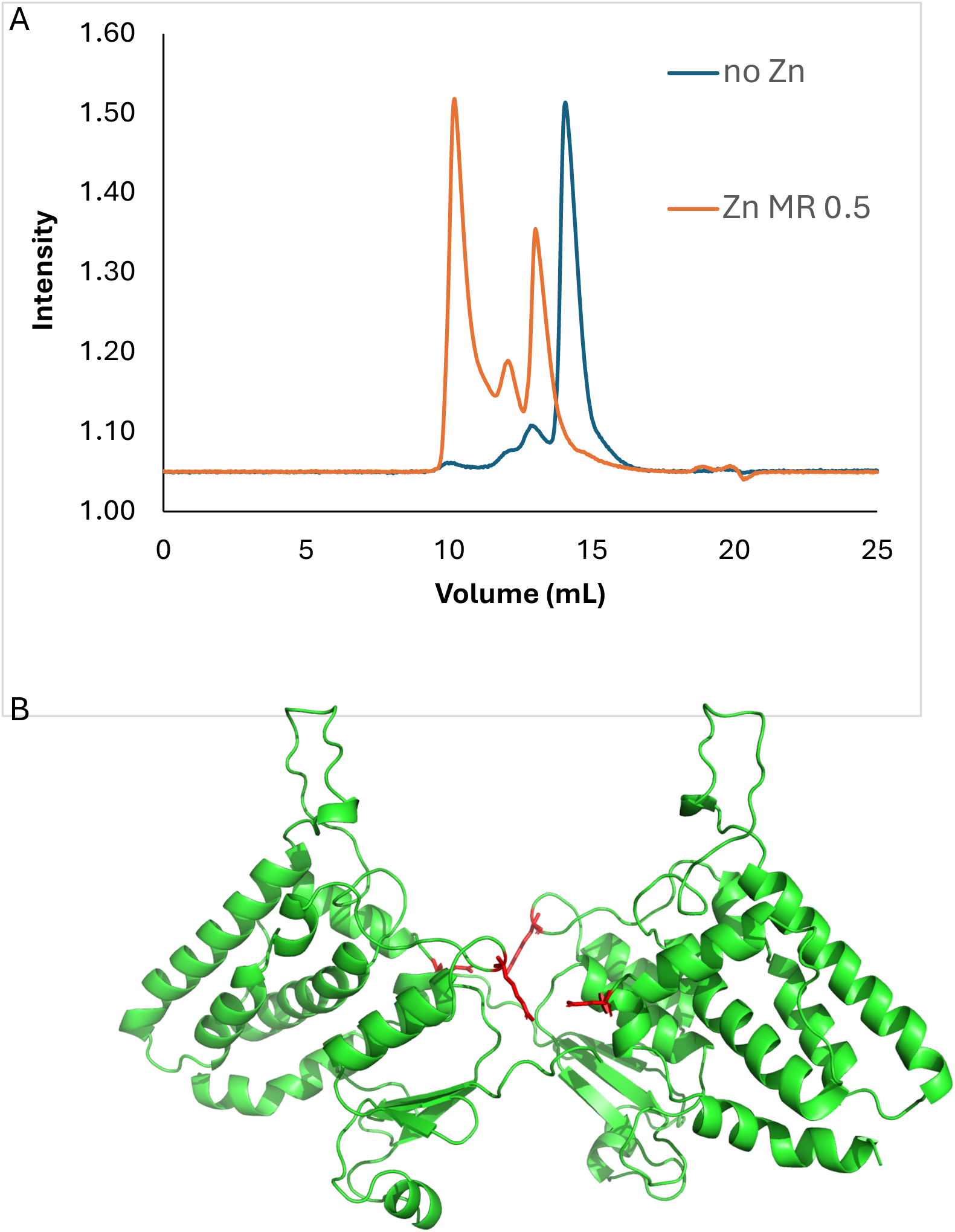
CLIC1 oligomeric equilibrium. A – SEC traces of CLIC1 in the presence (orange) and absence (blue) of a Zn^2+^:CLIC1 molar ratio of 0.5. B – SAXS-validated Alphafold multimer structural model of CLIC1 dimer. Key interactions between the intradomain loop and Helix_216-223_ are highlighted in red.

Alphafold multimer was used to predict the CLIC1 dimer structure (Figure 2B). This structure accurately matches the SAXS scattering data obtained for the dimer form. The predicted dimer structure suggests that key interactions holding the dimer interface involve the N-terminal domain, as well as the interdomain loop and the helix between residues 216-223. This is consistent with the data indicating an N terminal tag interference with oligomerisation. Interestingly, when the experiment was repeated in the presence of 0.5MR Zn^2+^ the experimental Zn^2+^ -bound SAXS data of the dimer no longer provided a good fit to the alpha fold dimer, suggesting that Zn^2+^ induces large conformational changes in CLIC1.

Mass photometry experiments on CLIC1 embedded on lipid nanodiscs revealed a tetrameric assembly, confirming that oligomerisation is required for CLIC1 membrane insertion (Figure S6).

### Zn^2+^ Triggers induces changes in dynamics and Oligomerization

To understand the effect of Zn^2+^ on CLIC1 structure and dynamics, titrations were followed by NMR. Zn^2+^ binding induced substantial chemical shift perturbations in the N-terminal domain and TM-adjacent regions (Figure 3A,B and S7). Residues 20–24, 65– 76, and 220–233 exhibited the most significant shifts, with the indole group of Trp35 splitting into two peaks, indicative of the presence of a second conformation induced by Zn^2+^.

**Figure 3.**
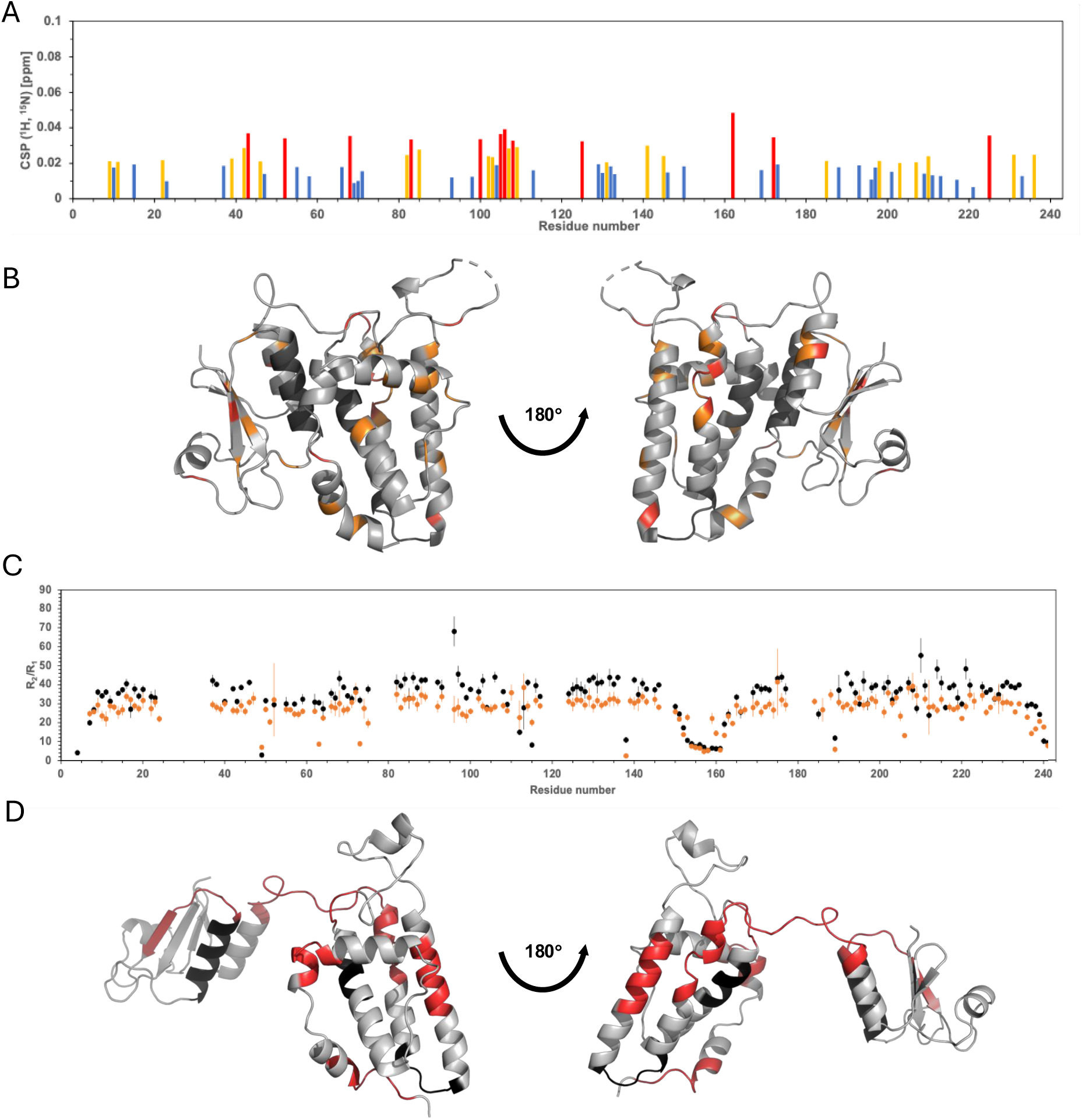
Zn^2+^-induced structural and dynamic changes. A – Chemical shift perturbation analysis of CLIC1 upon addition of Zn^2+^ to a molar ratio of 1. Residues showing large chemical shift perturbations (csp) are coloured red, medium csp in orange and non-signiFicant csp in blue. B – CLIC1 monomeric X-Ray structure (PDB 1k0n) showing areas affected upon Zn^2+^ binding. Regions with high csp are colour in red and medium csp in orange. Unassigned amino-acids resonances are coloured in black. C – R2/R1 values in the presence (orange) and absence (black) of a Zn^2+^:CLIC1 molar ratio of 0.5 obtained at 600 MHz. D – CLIC1 extended structure showing areas with enhanced dynamics upon Zn^2+^ binding coloured in red.

To avoid the contribution of high order oligomeric species of CLIC1, the monomeric N-terminal His-tag construct of CLIC1 was used to obtain relaxation rates in the presence of Zn^2+^. These revealed increased flexibility in residues across multiple residues in the N- and C-terminal domains and the interdomain loop, suggesting that Zn^2+^ promotes increased CLIC1 conformational heterogeneity, critical for membrane interaction (Figure 3C,D).

### Mutagenesis of residues in the putative TM alter CLIC1 insertion properties

Residues in CLIC1 TM, including R29 and W35, have been found to be involved in CLIC1 chloride channel properties. Alanine mutation of both residues showed significant differences in oligomerisation. The R29A mutation shifts the oligomerisation equilibrium towards a monomer, while the W35A mutation displays the opposite effect, promoting the formation of higher order oligomers (Figure 4A).

**Figure 4.**
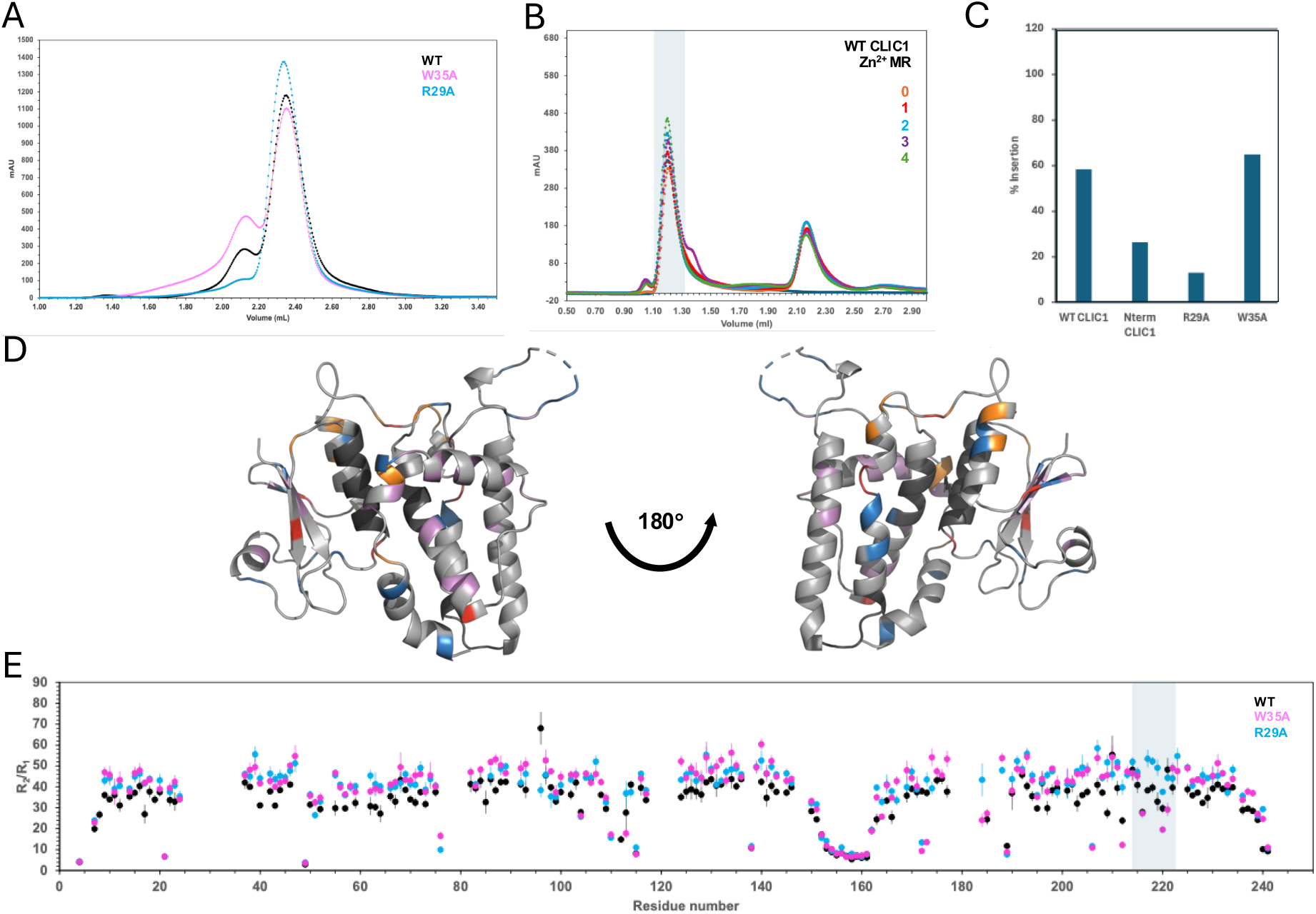
Impact of TM mutations on CLIC1 properties. A – SEC traces of CLIC1 (black), the W35A mutant (magenta) and the R29A mutant (blue), indicating differences in the relative populations of monomer (Elution volume of 2.35 mL) and dimer (Elution volume of 2.10 mL). B – SEC traces of insertion assay carried out in CLIC1 at increasing Zn^2+^ concentrations (from Zn^2+^:CLIC1 molar ratio 0 to 4). The vesicle peak is highlighted in grey. C – Insertion propensities of WT CLIC1, uncleaved N-terminal His-tagged CLIC1, R29A and W35A mutants, calculated from the intensity of the vesicle peak in the insertion assay. D – Chemical shift perturbations (csp) resulting from the R29A mutation. Regions with high csp are colour in red and medium csp in orange. Regions with resonances showing signiFicantly higher intensities compared to the WT are coloured pink, and those with signiFicantly lowre intensities compared to the WT are coloured blue. E – R2/R1 values of WT (black), the W35A mutant (magenta) and the R29A mutant (blue), obtained at 600 MHz.

An insertion assay was developed to quantify the level of membrane insertion of CLIC1. This assay involves incubation of CLIC1, Zn^2+^ and vesicles and injection in a SEC column which produces a SEC trace containing both a vesicle and a protein peak. The insertion of CLIC1 in vesicles at increasing Zn^2+^ concentrations was monitored through changes to the UV absorbance at the vesicle elution volume, which saw up to a 60% increase in CLIC1-vesicle association at increasing Zn^2+^ concentrations (Figure 4B and S8). Again, both mutations showed opposite effect, with R29A halting CLIC1 membrane insertion. W35A mutation enhanced CLIC1 membrane insertion in the absence of Zn^2+^ compared to the WT (Figure S7). The impact of both mutations on CLIC1 structure and dynamics was assessed by NMR. While significant differences were found between their ^15^N HSQC spectrum and the WT across the structure, mostly at the interface between the N- and C-terminal domains (Figure 4D), the changes in dynamics are localised around the helix between residues 216-223 (Figure 4E). The R29A mutation increases R2/R1 values in this helix, indicating rigidification, while W35A shows line broadening and lower R2/R1 values, and indication of enhanced dynamics.

## DISCUSSION

This study elucidates the intricate relationship between structural flexibility, oligomerization, and membrane insertion of CLIC1, providing significant insights into the role of dynamics and Zn^2+^ in regulating its function.

### Structural Flexibility as a Prerequisite for Membrane Insertion

Our findings establish that CLIC1’s metamorphic behaviour stems from its intrinsic structural flexibility. The importance of protein dynamics in CLIC membrane insertion has already been highlighted by previous studies, although lacking the atomic resolution needed to relate it to CLIC1 insertion mechanism(Ferofontov et al., 2018; Hossain et al., 2017; Stoychev et al., 2009). The separation of the N- and C-terminal domains in the minor elongated state exposes the TM region, enabling interaction with the lipid bilayer. SAXS and NMR data collectively highlight the importance of dynamic loops in facilitating these conformational changes. Previous FRET studies showed increased distance between residues in the N- and C-terminus domains in the membrane bound state (Goodchild et al., 2011). Comparative SAXS analysis of CLIC1 and GST O-1 (Figure S4), a structural homologue with no membrane-insertion properties, revealed that the elongated state is unique to the CLIC family, suggesting a specialized role in their metamorphic function.

### Zn^2+^ as a Trigger for Structural Rearrangement and Oligomerization

CLIC1 membrane insertion has been found to be triggered by Zn^2+^ binding (Varela et al., 2022). Zn^2+^ binding induces localized structural changes. The most significant shifts were observed in residues around the putative TM, and in residues 220–233, suggesting these regions play a central role in mediating the interaction with Zn^2+^. Relaxation rate measurements of monomeric CLIC1 in the presence of Zn^2+^ further corroborate the role of Zn^2+^ in modulating the protein’s dynamics. The increased flexibility observed suggests that Zn^2+^ binding destabilizes CLIC1 structural regions, both. This enhanced conformational heterogeneity facilitates the structural rearrangements necessary for membrane interaction, consistent with the hypothesis that Zn^2+^ acts as a modulator of CLIC1 membrane association. Previous research already established the role of divalent cations at triggering conformational changes and structure destabilisation to induce membrane association (Billen et al., 2008; Vasquez-Montes et al., 2019, 2022).

The oligomeric state of CLIC1 emerged as a critical determinant of its functional role, with the protein existing in a dynamic equilibrium between monomeric, dimeric, and higher-order oligomeric forms. SEC experiments revealed that the monomer is the predominant species under basal conditions, but the addition of Zn^2+^ shifted this equilibrium towards dimer and tetramer formation. The observed Zn^2+^-dependent oligomerization strongly correlates with increased membrane association, and correlates with the tetrameric CLIC1 form observed in lipid nanodiscs. The presence of an N-terminal tag inhibits both oligomerisation and the presence of elongated states, resulting in a significantly lower membrane insertion propensity (Figure 4C), highlighting the importance of this region in the overall dynamics and oligomerisation.

Mutagenesis of residues R29 and W35 provided further insights into the molecular determinants of CLIC1 membrane insertion. The R29A mutation disrupted oligomer formation, favouring the monomeric state, and abolished membrane insertion. NMR analysis revealed the rigidification of the helix spanning residues 216-223. Conversely, the W35A mutation, which promoted membrane insertion, was found to enhance higher-order oligomerization and enhance the flexibility of helix_216-233_. The structural model of CLIC1 dimer highlights key interactions at the dimer interface between protein N-terminus involving this helix, in a mechanism similar to other membrane-associated proteins(Quarato et al., 2016). Our findings suggest that the balance between rigidity and flexibility at the helix_216-233_ is crucial for CLIC1’s functional transitions and that Zn^2+^-mediated modulation of these properties is a key regulatory mechanism.

In conclusion, we provide a comprehensive structural and mechanistic model for CLIC1 transition from a soluble globular protein to a membrane-bound chloride ion channel (Figure 5). The identification of specific residues involved in oligomer stabilization further highlights potential targets for modulating CLIC1 activity therapeutically.

**Figure 5.**
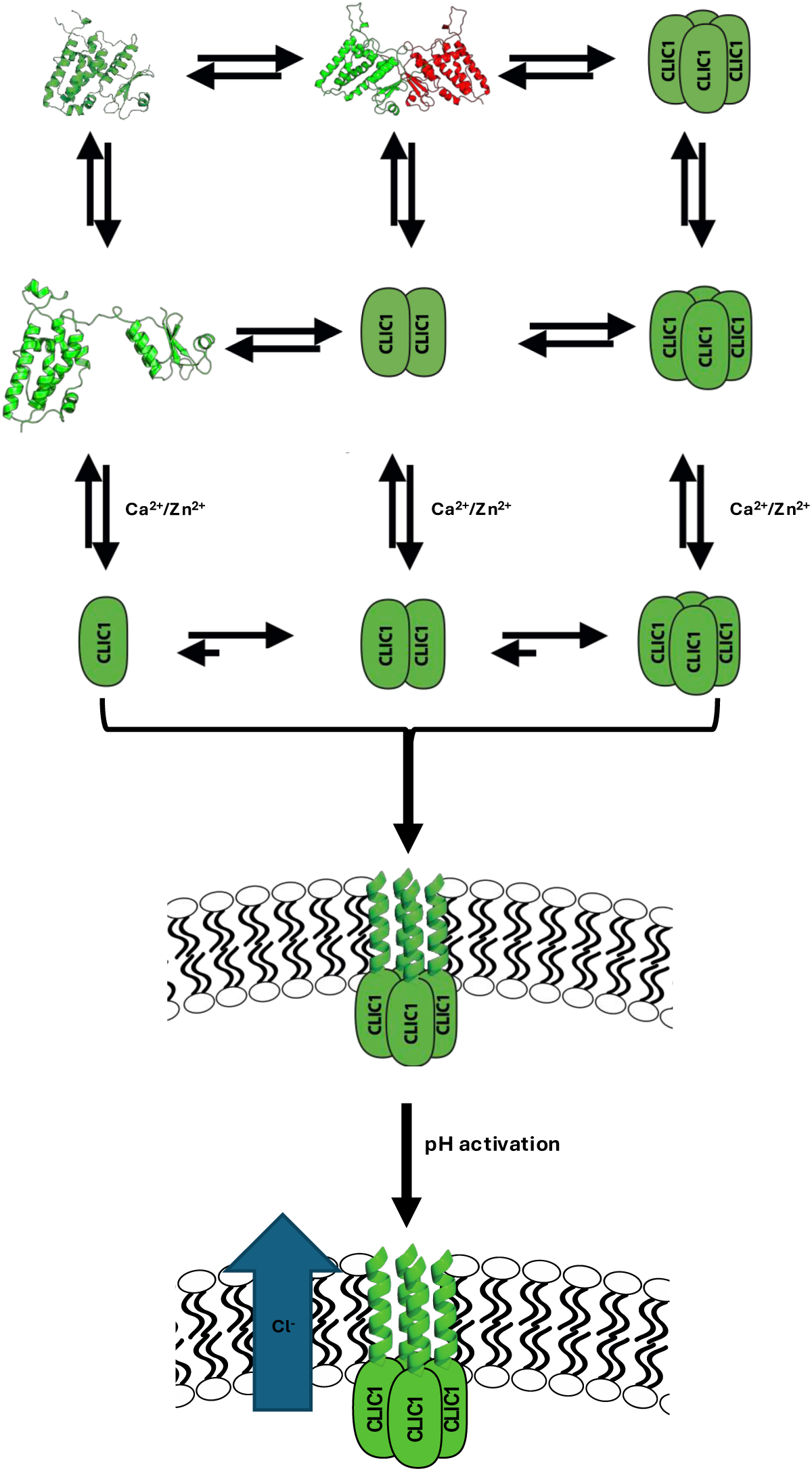
The Mechanism of CLIC1 membrane insertion. CLIC1 exists in solution as an equilibrium between monomer, dimer and tetrameric forms in a closed and an open state. Upon Zn^2+^, a destabilisation of these structures is produced, unravelling structural changes on the N-terminal domain and promoting oligomerisation, which in the presence of a lipid bilayer leads to membrane insertion into a tetrameric assembly. Chloride channel activation is activated upon acidiFication. Green elongated shapes are used for those species whose structure is still unknown.

## ACKNOWLEDGEMENTS

We acknowledge support from the Wellcome Trust Seed Award (207743/Z/17/Z) and funding from the Alan Turing Institute. Joseph Cassar was supported by a PhD studentship awarded by the University of Kent. This work was supported by the Francis Crick Institute through provision of access to the MRC Biomedical NMR Centre. The Francis Crick Institute receives its core funding from Cancer Research UK (CC1078), the UK Medical Research Council (CC1078), and the Wellcome Trust (CC1078).

