## Supplementary figures for "Mechanistic Elucidation of CLIC1 Membrane Insertion via Structural and Dynamic Modulation"

### SUPPLEMENTARY INFORMATION

FIGURE S1

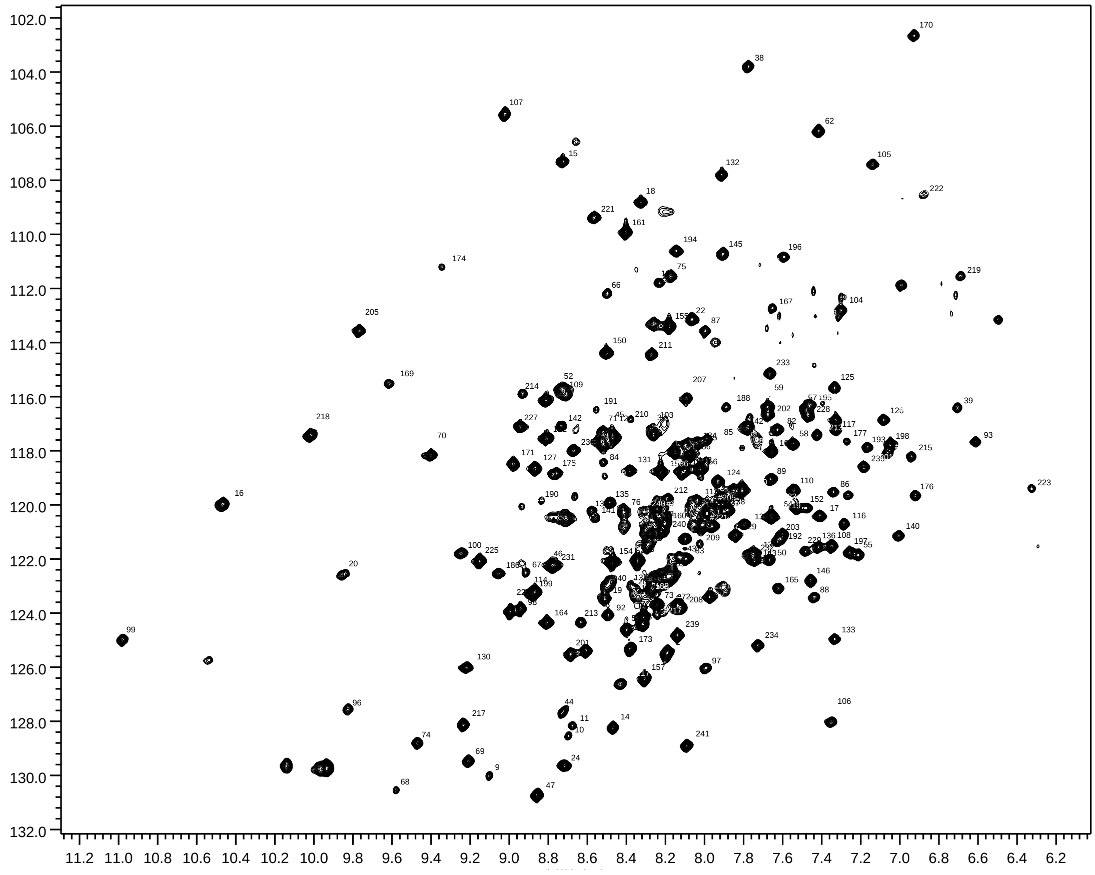


**Figure S1. Assigned $^{15}N$ TROSY HSQC spectrum of CLIC1**

FIGURE S2

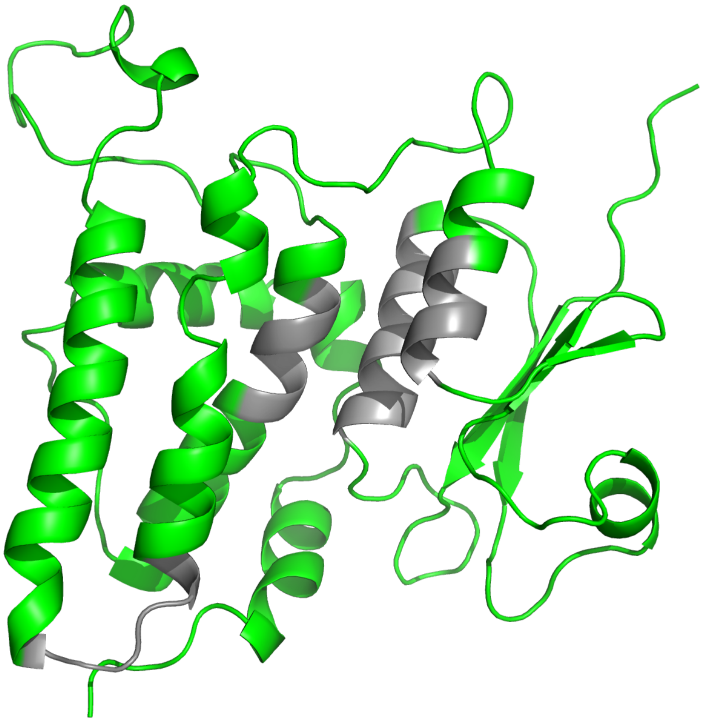

**Figure S2. CLIC1 backbone assignment coverage.** CLIC1 monomeric X-Ray structure (PDB 1k0n) showing areas with (green) and without (grey) assigned backbone resonances.

FIGURE S3

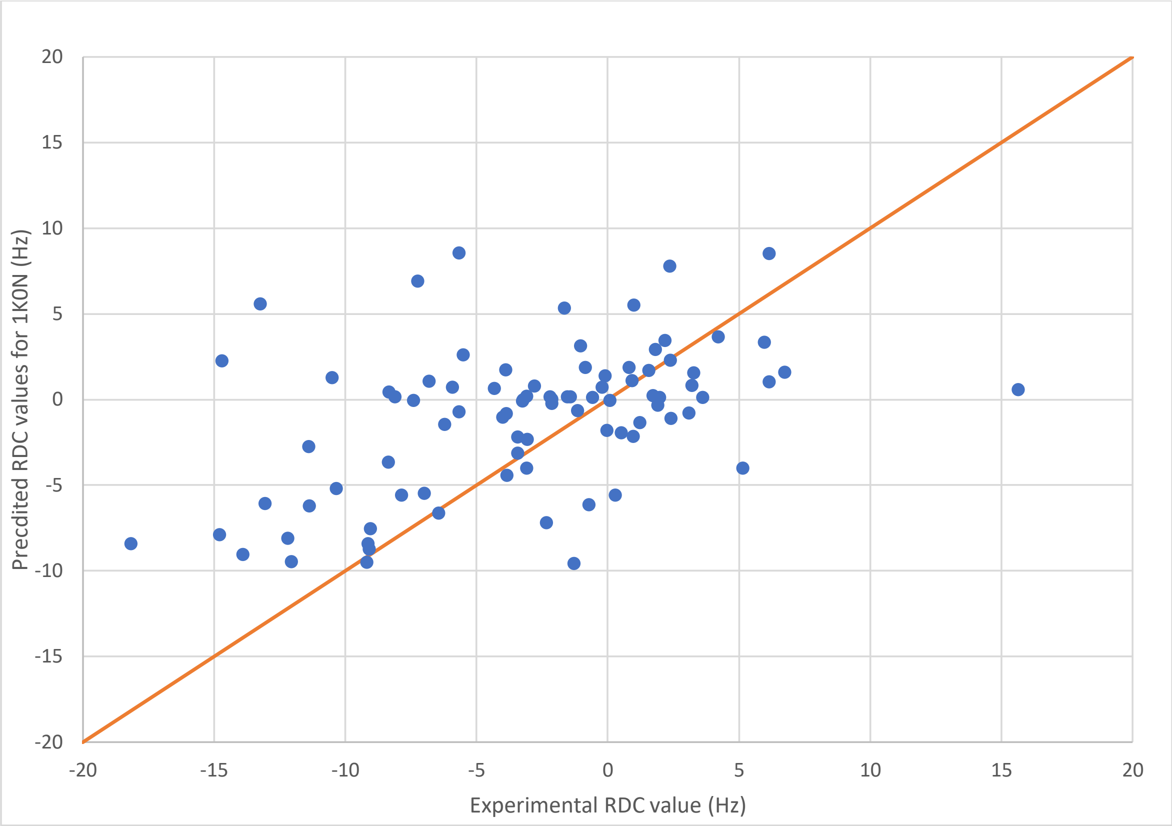


**Figure S3. Differences between X-Ray and solution structures of CLIC1.** NH experimental versus predicted RDCs from CLIC1 X-Ray structure 1k0n showing poor agreement.

FIGURE S4

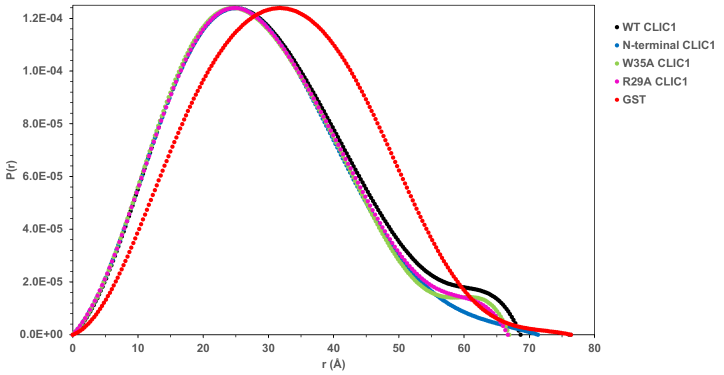


**Figure S4. SAXS shows different average shapes for CLIC1 and omega GST in solution.** SAXS Pair distribution function ($P_R$) of CLIC1 demonstrates CLIC1 is globular with an extended tail that suggests an elongated conformation. $P_R$ plots for WT CLIC1 (black), N-terminal His-tagged CLIC1 (blue), the R29A (magenta) and W35A (green) mutants and omega-1 GST (red) shows differences in the contribution of this elongated conformation.

FIGURE S5

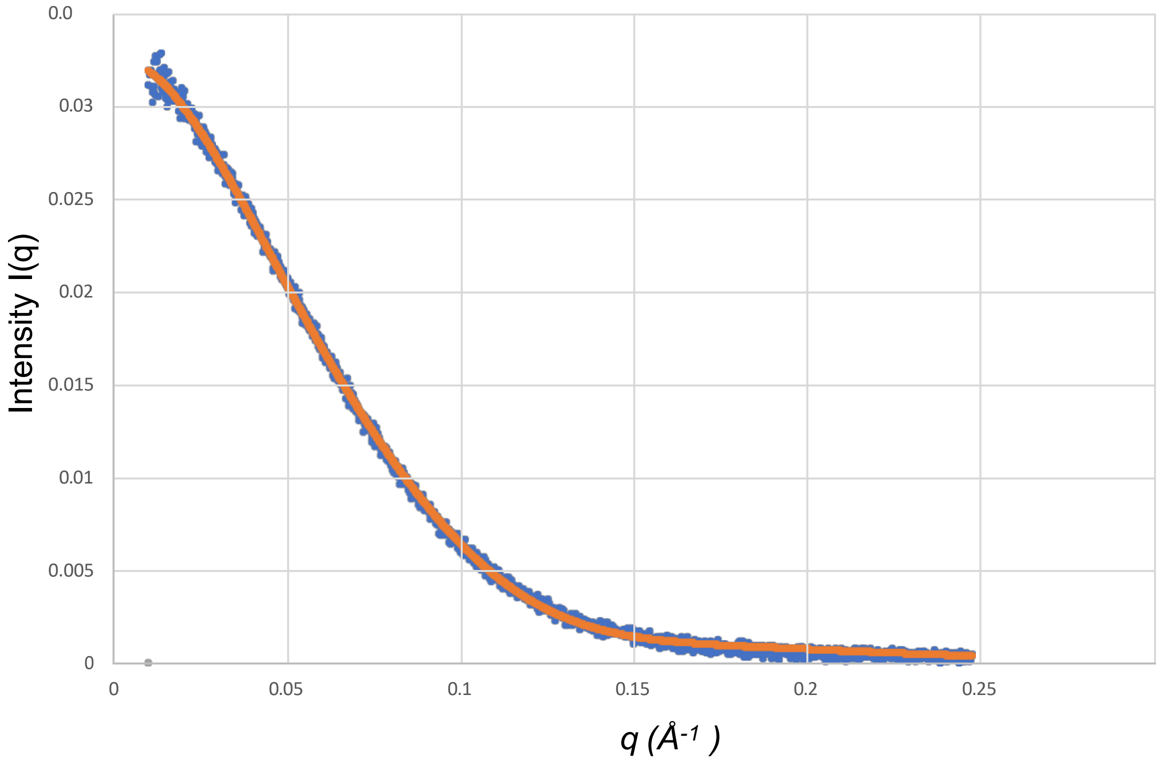


**Figure S5. Comparison of the EOM ensemble of CLIC1 structures against the experimental scattering data**. EOM ensemble was calculated as a weighted profile and then plotted in orange whilst the experimental scattering data for CLIC1 is shown in blue. The ensemble of structures provides a good fit to the experimental data giving a chi squared score of 1.04.

FIGURE S6

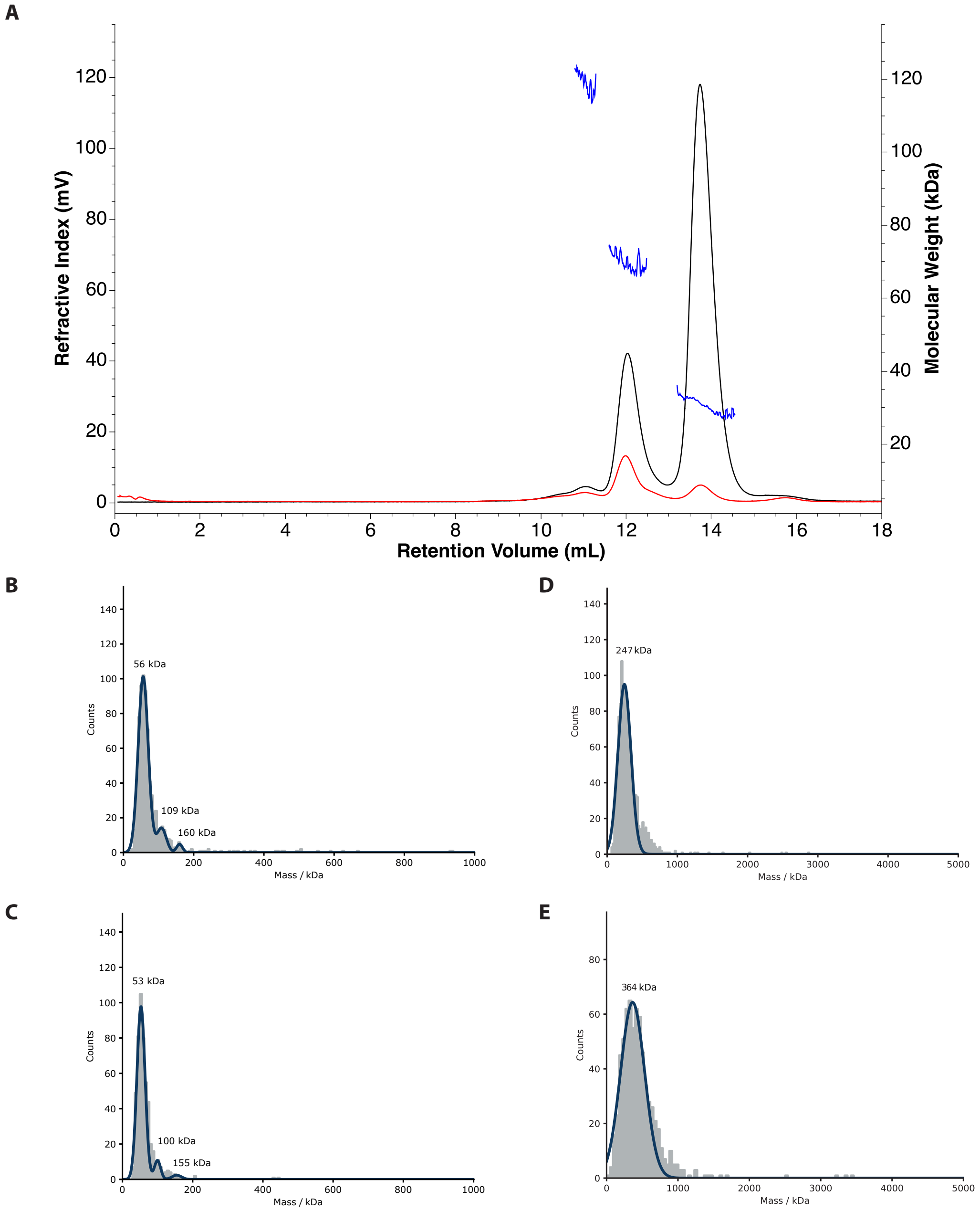


**Figure S6. CLIC1 oligomeric states in solution and in the chloride channel form.** SEC-MALS traces of non-treated CLIC1 (black). CLIC1 samples corresponding to the dimeric form were subjected for a second SEC-MALS run (red). The molecular weights calculated for the monomer, dimer and tetramer peaks are shown in blue. B,C – Mass Photometry histograms for 100nM CLIC1 samples in the absence and presence of an equimolecular concentration of $Zn^{2+}$. D,E – Mass Photometry histograms of empty and CLIC1-containing nanodiscs.

FIGURE S7

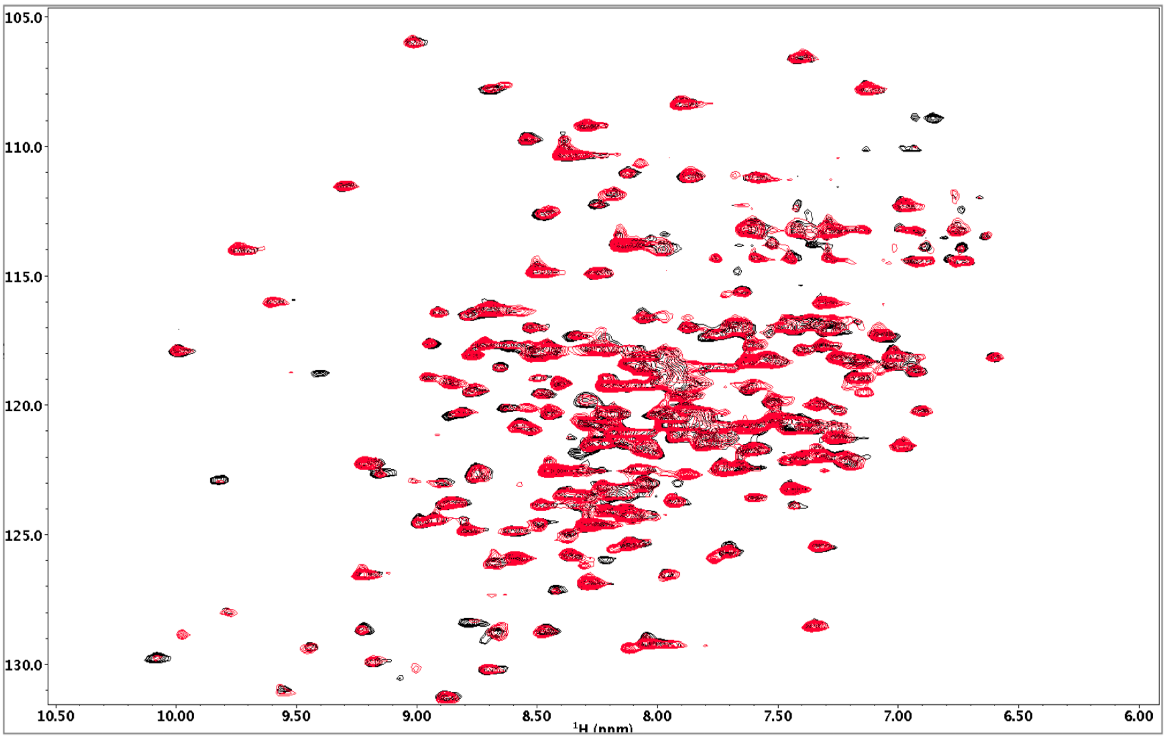


**Figure S7. NMR $Zn^{2+}$ titration.** $^{15}N$ TROSY HSQC spectra of CLIC1 in the absence (black) and presence (red) of $ZN^{2+}$ in a $Zn^{2+}$:CLIC1 molar ratio of 1.

FIGURE S8

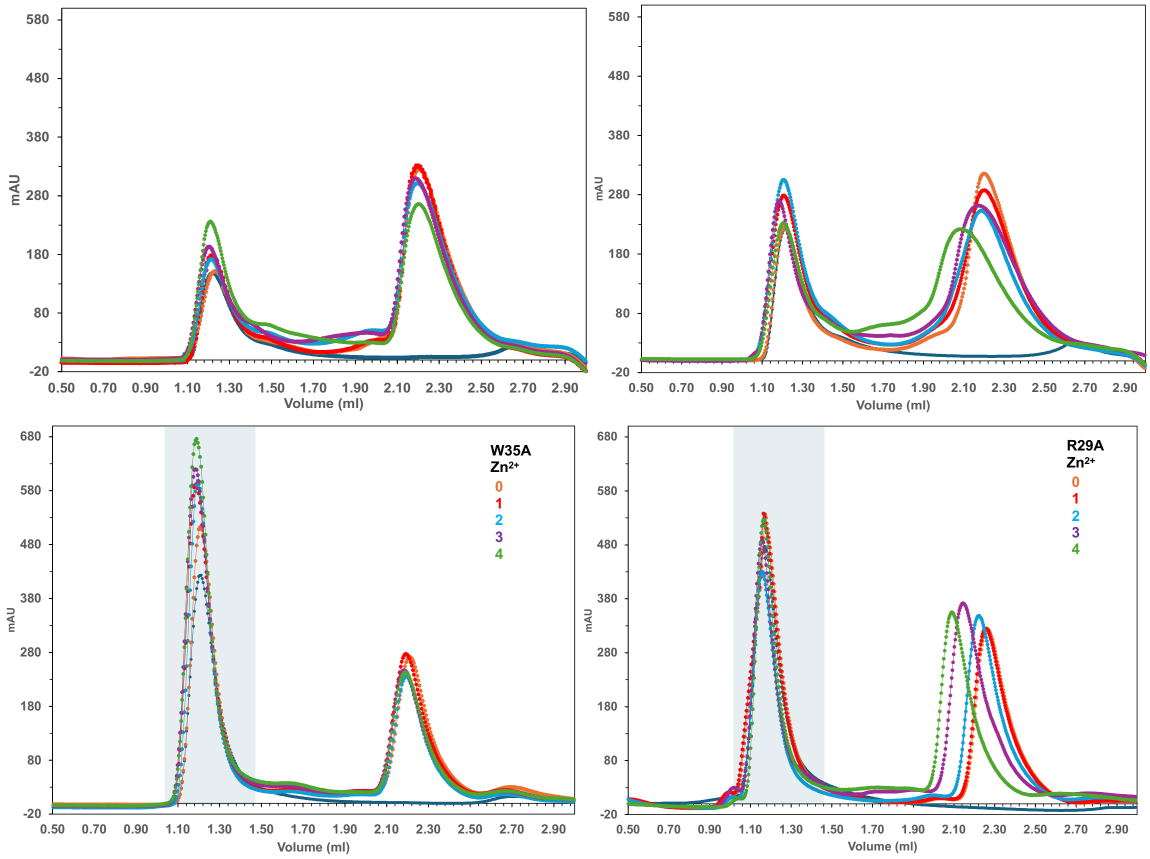
